# Assembly of a high-quality reference genome for the rat tapeworm *Hymenolepis diminuta*

**DOI:** 10.64898/2026.06.23.734100

**Authors:** Saurav K. Choudhary, Nikitha Sundaresha, Kaixiong Ye, Casey M. Bergman, Tania Rozario

## Abstract

The rat tapeworm, *Hymenolepis diminuta*, is an important laboratory model for uncovering molecular processes that underly the success of tapeworms as parasites. Despite its importance, a high-quality reference genome for this species is lacking. Here we present a highly contiguous and effectively complete genome of *H. diminuta* assembled from PacBio HiFi long-read sequencing data. Our primary assembly consists of 7 scaffolds (N50=29.25 Mb) with total length of 186.53 Mb, has only 7 gaps, and contains 95.7% complete Lophotrochozoan BUSCOs. Our assembly allows us to confirm aspects of *Hymenolepis* genome organization, such as high repeat content and unusual chromosomal ends, and to show that *Hymenolepis* genomes encode ∼10,000 genes. Together with annotations of nuclear tRNAs, mtDNA protein coding genes, and mtDNA tRNAs, our assembly currently provides one of the most complete genome resources for a tapeworm species and will enable research on parasitism, animal regeneration, development, and evolution.

## Introduction

Flatworms (Platyhelminthes) include both free-living and parasitic taxa, the latter forming a monophyletic clade (Neodermata) that includes all flukes and tapeworms (Collins 2017). Parasitic flatworms infect humans, livestock, and companion animals, causing symptoms such as malnourishment, morbidity, intestinal distress, anemia, organ damage/failure, seizures, certain cancers, and even death (Craig and Ito 2007; McManus et al. 2018; Brindley et al. 2021; Kusnoto et al. 2025). Improved genomic resources are needed to better understand parasite biology and support diagnostic and therapeutic strategies. However, studies of tapeworms are often impeded by their complex life cycles and host ranges.

The rat tapeworm, *Hymenolepis diminuta*, is an established experimental model because of several key advantages. Its life cycle is easily maintained using rats and beetles, functional studies can be performed using RNA interference, and both adult and larval stages can be cultured axenically. In addition, numerous staining methods are available to examine gene expression and tissue organization (Evans 1980; Sukhdeo and Sukhdeo 1994; Rozario and Newmark 2015; Rozario et al. 2019; Ishan et al. 2025; Nanista et al. 2025). Despite these attributes, a high-quality genome assembly and gene annotation for *H. diminuta* is lacking.

An initial draft genome assembly for *H. diminuta* was generated as part of a wider effort to sequence over 100 parasitic helminth genomes (International Helminth Genome Consortium 2019). A subsequent hybrid short- and long-read assembly improved contiguity but remained highly fragmented with 719 scaffolds (Nowak et al. 2019). This exceeds the haploid chromosome number (n=6) reported for this species (Jones 1945; Kisner 1957; Mutafova and Gergova 1994). Nowak et al. (2019) predicted 15,169 genes, which is substantially higher than the 10,139 genes annotated in a closely related species, *Hymenolepis microstoma* (Olson et al. 2020). Because *H. microstoma* has one of the most complete tapeworm genome assemblies currently available (Kamenetzky et al. 2022), the elevated gene count in *H. diminuta* is likely inflated.

Recent advances in long-read sequencing and assembly have enabled chromosome-scale genome assemblies. Olson et al. (2020) assembled the nuclear genome for the tapeworm *H. microstoma* into six scaffolds corresponding to the haploid chromosome complement, with minimal gaps. This assembly revealed atypical chromosomes that terminate with either telomeric or centromeric arrays, leading to the proposal that centromeric arrays have replaced previously existing telomeric arrays at one end of each chromosome (Olson et al. 2020). This differs from the canonical arrangement of telomeres at both chromosomal ends in other tapeworm species (Tsai et al. 2013; Li et al. 2018). Whether this unusual chromosome organization is specific to *H. microstoma* or found in other tapeworms remains unresolved.

Here we report a highly contiguous and effectively complete genome assembly for *H. diminuta* generated using PacBio long-read sequencing. This assembly is a substantial improvement relative to previous versions and enables reassessment of gene number and genome organization in this species. Our assembly and annotation provide a resource for comparative genomics and future genome-based analyses such as single-cell and spatial transcriptomics, gene regulatory network predictions, and chromatin-level investigations, thus expanding the utility of this important model system.

## Methods and Materials

### Animal Rearing and Care

We generated a genetically homogeneous line of *H. diminuta* originally obtained from Carolina Biological (132232) by inbreeding for 15 generations as follows. A single Sprague-Dawley rat was infected with a single cysticercoid that was grown to reproductive maturity, at which time the mature proglottids were used to infect a batch of 50 mealworm beetles (*Tenebrio molitor*) from which a single cysticercoid was selected, and the process repeated. To infect rats, cysticercoids were fed by oral gavage in ∼0.5 mL of 0.85% NaCl and reared until the tapeworms reached gravidity. Rats were euthanized in a CO2 chamber and tapeworms were flushed from the small intestine into 1X Hanks Balanced Salt Solution (HBSS; Corning) (140 mg/L CaCl2, 100 mg/L MgCl2.6H2O, 100 mg/L MgSO4.7H2O, 400 mg/L KCl, 60 mg/L KH2PO4, 350 mg/L NaHCO3, 8 g/L NaCl, 48 mg/L Na2HPO4, 1 g/L D-glucose, no phenol red). To infect beetles, ∼10 cm of the most mature gravid proglottids were smashed with apple sauce onto filter paper and fed to beetles that were pre-starved for 3-5 days. Rodent care was performed in accordance with protocols approved by the Institutional Animal Care and Use Committee of the University of Georgia (A2023 10-019-A8).

### DNA extraction, PacBio Library Preparation, and Genome Sequencing

980 mg of roughly dried tissue from 6-day-old *H. diminuta* adults was frozen on dry ice and sent to the Arizona Genomics Institute for DNA extraction, library preparation, and PacBio HiFi sequencing. High molecular weight (HMW) DNA was extracted from homogenized tissue in Tris HCl buffer 0.1M pH 8.0, EDTA 0.1M pH8, SDS 1% and Proteinase K at 50°C for 60 minutes. The mixture was spun down, and the aqueous phase was transferred to a new tube. 5M Potassium acetate was added, precipitated on ice, and spun down. After centrifugation, the supernatant was gently extracted with 24:1 chloroform:isoamyl alcohol. The upper phase was transferred to a new tube and DNA precipitated with isopropanol. DNA was collected by centrifugation, washed with 70% ethanol, air dried, and dissolved thoroughly in 10 mM Tris-HCl followed by RNAse treatment. DNA purity was measured with Nanodrop, DNA concentration measured with Qubit HS kit (Invitrogen), and DNA size was validated by Femto Pulse System (Agilent). Extracted HMW DNA was sheared to an appropriate size range (10-20 kb) using Megaruptor 3 (Diagenode) followed by SMRTbell cleanup beads. The sequencing library was constructed following manufacturer’s protocols using SMRTbell Prep kit 3.0. The final library was size selected on a Pippin HT (Sage Science) using S1 marker with a 10-25 kb size selection. The recovered final library was quantified with Qubit HS kit (Invitrogen) and size checked on Femto Pulse System (Agilent). The final library was prepared for sequencing with PacBio Sequel II Sequencing kit 3.1 for HiFi library, loaded on a single Revio SMRT cell, and sequenced in CCS mode for 24 hours.

### Genome Assembly and Quality Assessment

PacBio HiFi reads were used to estimate genome size, repetitiveness, and heterozygosity with GenomeScope (v2.0; p=2 k=21) (Ranallo-Benavidez et al. 2020) based on a kmer histogram generated by meryl (v1.4.1; k=21) (Rhie et al. 2020). HiFi reads were then assembled using hifiasm (v0.19.40; default settings) (Cheng et al. 2021). The primary hifiasm assembly was scaffolded with RagTag (v2.1.0; default settings) (Alonge et al. 2022) using the most closely related species with a chromosome-level genome assembly, *H. microstoma* (Genbank: GCA_000469805.3) (Olson et al. 2020).

We were unable to identify a complete *H. diminuta* mtDNA contig in the scaffolded hifiasm assembly, however we did identify two contigs that partially matched the *H. microstoma* mtDNA. These two partial mtDNA contigs were removed from the scaffolded assembly and a complete mtDNA genome was assembled from raw PacBio HiFi reads using MitoHiFi (v3.2.1; default settings) (Uliano-Silva et al. 2023) guided by a previously reported *H. diminuta* mtDNA genome (Genbank: NC_002767) (von Nickisch-Rosenegk et al. 2001). The resulting complete mtDNA genome was reoriented to start with the *cox1* gene using seqkit (v2.9.0) (Shen et al. 2024), then appended to the scaffolded nuclear assembly to create the final “chromosome-level” assembly used for most analyses in this report (unless otherwise noted). An additional 191 unplaced contigs were appended to the chromosome-level assembly to generate the final assembly that was deposited at DDBJ/ENA/GenBank under the accession JBVFLO000000000.

Assembly statistics were computed using GFAstats (v1.3.11; --stats, --segment-report, --path-report) (Formenti et al. 2022). Assembly completeness was assessed using BUSCO (5.8.3; default settings) (Manni et al. 2021) with the lophotrochozoa_odb12 database (Tegenfeldt et al. 2024). Correctness of our final submitted assembly was assessed using Inspector (v1.3.1; --datatype hifi) (Chen et al. 2021).

### Repeat Identification and Masking

We identified repetitive element families in our *H. diminuta* chromosome-level assembly using RepeatModeler (v2.0.3; -LTRStruct) (Flynn et al. 2020). The resulting library of repeats was then used as an input for RepeatMasker (v4.1.4; -xsmall) (http://www.repeatmasker.org) to annotate and softmask repetitive elements in the *H. diminuta* genome. Similar procedures were applied to the *H. microstoma* genome from Olson et al. (2020) (Genbank: GCA_000469805.3) to allow controlled comparison of interspersed repeat content between the two species. The RepeatModeler library generated from our *H. diminuta* assembly was also used to annotate repeats in the Nowak et al. (2019) assembly (Genbank: GCA_902177915.1) using RepeatMasker (v4.1.4; -xsmall) to allow controlled comparison of interspersed repeat content between the two *H. diminuta* assemblies.

### Genome Annotation and Quality Assessment

Gene prediction was performed with BRAKER3 (v3.0.3, --gff3) (Gabriel et al. 2024), under three evidence configurations: (i) protein-only, (ii) RNA-seq–only, and (iii) combined RNA-seq + protein evidence. To evaluate potential run-to-run variability in our genome annotation process, each genome-by-evidence combination was replicated five times. For all BRAKER3 runs, we used the soft-masked versions of our chromosome-level *H. diminuta* assembly, the *H. diminuta* from Nowak *et al*. (2019), and the *H. microstoma* from Olson *et al*. (2020) described above. BRAKER3 runs based on protein evidence used the OrthoDB (v12) Metazoa dataset (Tegenfeldt et al. 2024). BRAKER3 runs based on RNA-seq evidence used BAM files generated using STAR (v2.7.10; parameters: --outSAMstrandField intronMotif, --outSAMtype BAM SortedByCoordinate) (Dobin et al. 2012) with the following inputs: SRA IDs SRR9295549-SRR9295553 and SRR9275651-SRR9275655 (Rozario et al. 2019) for both *H. diminuta* assemblies; and SRA IDs ERR225719-ERR225730, ERR337915, ERR337928, ERR337940, ERR337952, ERR337964, ERR337976 (Olson et al. 2020) for the *H. microstoma* assembly. For downstream analyses, we used the braker.gff3 files from each run, which contains transcript models jointly inferred by AUGUSTUS and GeneMarkS-T. Protein coding gene models predicted by BRAKER3 represent coding exons only and do not include untranslated regions.

For our final protein coding gene annotation, we selected a single BRAKER3 replicate for each genome that used combined RNA-seq + protein input. The chosen replicate was the one whose total predicted protein coding gene count was closest to the median across the five replicate runs. Functional annotation of the final predicted protein coding sequences set was performed using EggNOG-mapper (v2.1.9; -m diamond --decorate_gff --decorate_gff_ID_field ID) (Cantalapiedra et al. 2021) and the EggNOG (v5.0) database (Huerta-Cepas et al. 2019). EggNOG-based functional attributes were directly transferred onto the corresponding BRAKER3 GFF3 gene models using the ID field to link proteins and genomic features.

Transfer RNA (tRNA) genes were identified using tRNAscan-SE (v2.0.12; -E -j) (Chan et al. 2021) in eukaryotic mode using the soft-masked *H. diminuta* genome as input and output as GFF3. Our *H. diminuta* mitochondrial genome assembly was annotated using MitoFinder (v1.4.1; --new-genes -o 9) (Allio et al. 2020) using the previously published *H. diminuta* mtDNA (Genbank: NC_002767) as input. Nuclear tRNA gene and all mtDNA annotations were subsequently merged with the final functionally annotated BRAKER3 protein coding gene set to generate a GFF3 file representing the complete genome annotation. During post-annotation quality control analyses, we identified 123 nuclear gene models that were completely overlapped by other annotated genes. The shorter gene of fully overlapping gene model pairs was removed to eliminate redundancy and create a final GFF3 file that was compatible with NCBI genome submission requirements.

Quantification of genome annotation statistics was performed using the Genome Annotation Generator (GAG) (v2.0.1; default settings) (Geib et al. 2018). To ensure consistency with data reported in previous publications we used GFF3 files from Nowak *et al*. (2019), and Olson *et al*. (2020) obtained from WormBase ParaSite (release WBPS19). To directly compare gene models from different *H. diminuta* assemblies, we transferred coordinates of nuclear protein coding gene models from the Nowak *et al*. (2019) assembly onto our final assembly using Liftoff (v1.6.3; default settings) (Shumate and Salzberg 2021), excluding features mapping to unplaced contigs. We performed annotation comparisons in two ways: first using “all isoforms” per gene and second using only the “longest isoform” per gene. The longest isoform analysis attempts to control for differences in how alternative splice forms are annotated across studies and to simplify interpretation of annotation overlaps. The longest isoform per gene was extracted using agat_sp_keep_longest_isoform.pl (v1.4.2; -gff) (Dainat 2024). Both annotation comparisons were performed using agat_sp_compare_two_annotations.pl (v1.4.2; default settings) to assess gene overlaps.

We performed BUSCO analysis at the transcriptome level using the longest isoform per gene, which avoids false inference of duplicated BUSCO genes. Coding sequences corresponding to the longest isoforms were extracted using SeqKit (v2.8.2) (Shen et al. 2024) and used as input for BUSCO (v5.8.3; -m transcriptome) (Manni et al. 2021) to evaluate protein coding gene annotation completeness relative to the lophotrochozoa_odb12 database.

### Comparative genomics

Synteny analyses between *H. diminuta* and *H. microstoma* was performed using GENESPACE (v1.2.3; default parameters) (Lovell et al. 2022) using the longest isoform per gene. Orthogroups inferred from protein sequences were predicted using OrthoFinder (v2.5.5) (Emms and Kelly 2019) and supplied as input to GENESPACE to generate riparian plots. Syntenic blocks between *H. diminuta* and *H. microstoma* were identified using the syntenicBlock_coordinates.csv file generated by GENESPACE. Coordinates of putative inversion breakpoints were identified by filtering for syntenic blocks with a negative “orient” field, then identifying the intervals from the ends of inverted syntenic blocks to their neighboring 5’ and 3’ syntenic blocks, respectively. Assessment of assembly quality in putative breakpoint regions was assessed by filtering for high-quality mapped reads (MAPQ≥30) from BAM files generated by Inspector and visualizing evidence for contiguous overlapping read support in IGV (Robinson et al. 2011) in the context of assembly gaps, genes, and interspersed repeats. Whole genome alignment of the *H. diminuta* assembly versus the *H. microstoma* genome was performed using nucmer (v4.0.0; default settings), delta-filter (v4.0.0; -1), and mummerplot (v4.0.0; --size large --color -f --png) (Marçais et al. 2018).

### Telomere and Centromere Identification

Telomeric repeats in the *H. diminuta* genome were identified using Tidk search (v0.2.65; -s) (Brown et al. 2025) using the canonical telomeric repeat TTAGGG. Putative centromeric regions in the *H. diminuta* genome were identified using centroAnno (v1.0.2; -x anno-asm) (Qi et al. 2025). Centromere annotations were formatted using the cautils.py utility from the centroAnno distribution, and both telomere and centromere annotations were visualized using a custom python script.

## Results and Discussion

### PacBio HiFi long-reads can generate a highly contiguous and effectively complete tapeworm genome assembly

We generated >1.3 million PacBio Hifi reads (N50 length = 17.2 kb) from a population of highly-inbred adult *H. diminuta* tapeworms. Using kmer frequencies generated from unassembled HiFi reads, we estimated the length of *H. diminuta* genome to be 190.5 Mb, with a high repeat content (21.8%) and low level of heterozygosity (0.05%) (Supplementary Figure 1). We assembled the complete set of HiFi reads (∼110-fold coverage) with no additional input data using hifiasm (Cheng et al. 2021), which generated a primary assembly of 206 contigs in pseudo-haplotype format. Primary assembly contigs were scaffolded using the chromosome-level assembly of the mouse bile-duct tapeworm, *H. microstoma* (Olson et al. 2020). We were unable to identify a complete mtDNA genome in our initial hifiasm assembly, so we assembled the *H. diminuta* mtDNA from raw reads using a previously reported complete *H. diminuta* mtDNA genome as a reference (von Nickisch-Rosenegk et al. 2001). The resulting mtDNA assembly for our strain was added to the nuclear scaffolds to create a chromosome-level assembly of 7 scaffolds made up of 14 contigs, with scaffold and contig N50s of 29.2 Mb and 24.7 Mb, respectively (Table 1). These 7 chromosome-level scaffolds have a total length of 186.5 Mb, which corresponds to ∼98% of the genome size estimated from unassembled reads. Two unplaced contigs matching partial mtDNA sequences were excluded from the primary hifiasm assembly and the remaining 191 unplaced contigs (totaling 14.8 Mb) were appended to the chromosome-level assembly prior to submission to NCBI. We validated the integrity of our final assembly using Inspector, which found no evidence for any small-scale misassemblies relative to mapped read support. Unless otherwise noted, all analyses and results reported here use the chromosome-level assembly.

**Table 1.**
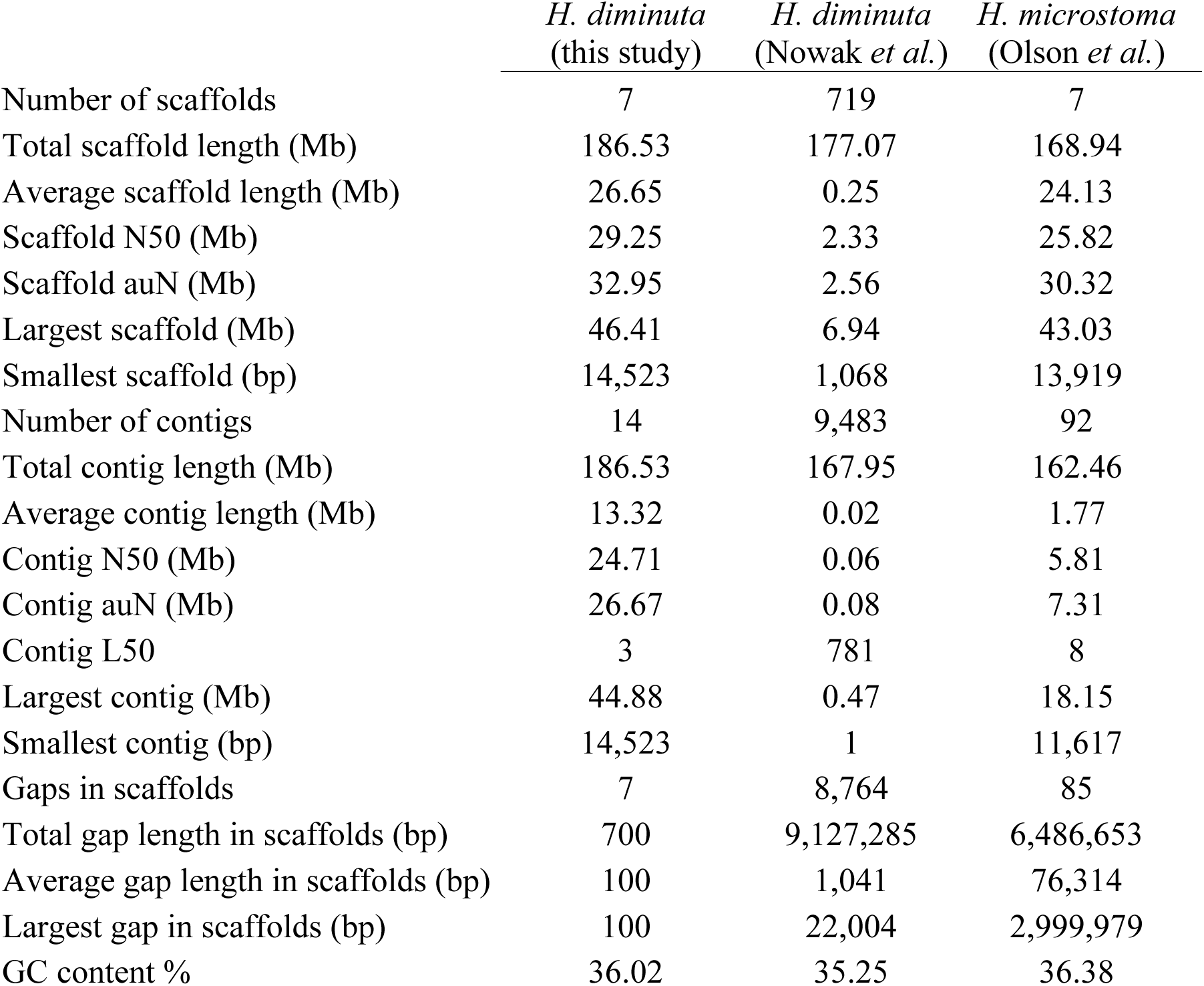
Assembly statistics for *Hymenolepis* genomes. Statistics shown are for our chromosome-level assembly of *H. diminuta* (excluding unplaced contigs), the *H. diminuta* assembly from Nowak et al. (2019) (Genbank: GCA_902177915.1) and the *H. microstoma* assembly from Olson et al. (2020) (Genbank: GCA_000469805.3).

Our HiFi-only chromosome-level assembly represents a substantial improvement in contiguity relative to the most recent scaffold-level version of the *H. diminuta* genome reported by Nowak et al. (2019) (Table 1). Compared to the hybrid assembly of Nowak et al. (2019), our assembly has >100-fold fewer scaffolds, >100-fold longer average scaffold length, and >1300-fold fewer gaps within scaffolds. Our *H. diminuta* assembly also exceeds contiguity metrics for the chromosome-level assembly of the congener *H. microstoma* (Olson et al. 2020), which is regarded as one of the best-assembled genomes in the class Cestoda (Kamenetzky et al. 2022). Notably, our assembly has >24 Mb of additional non-gap sequence, >7-fold larger average contig length, 10-fold fewer gaps than the Olson et al. (2020) *H. microstoma* assembly. Our chromosome-level assembly also has a very high level of estimated completeness based on the high proportion of complete BUSCO genes (95.7%) present in the Lophotrochozoan OrthoDB database (Tegenfeldt et al. 2024) (Figure 1). These results indicate that HiFi-based assembly of inbred tapeworms can lead to a highly contiguous and effectively complete genome assembly.

**Figure 1.**
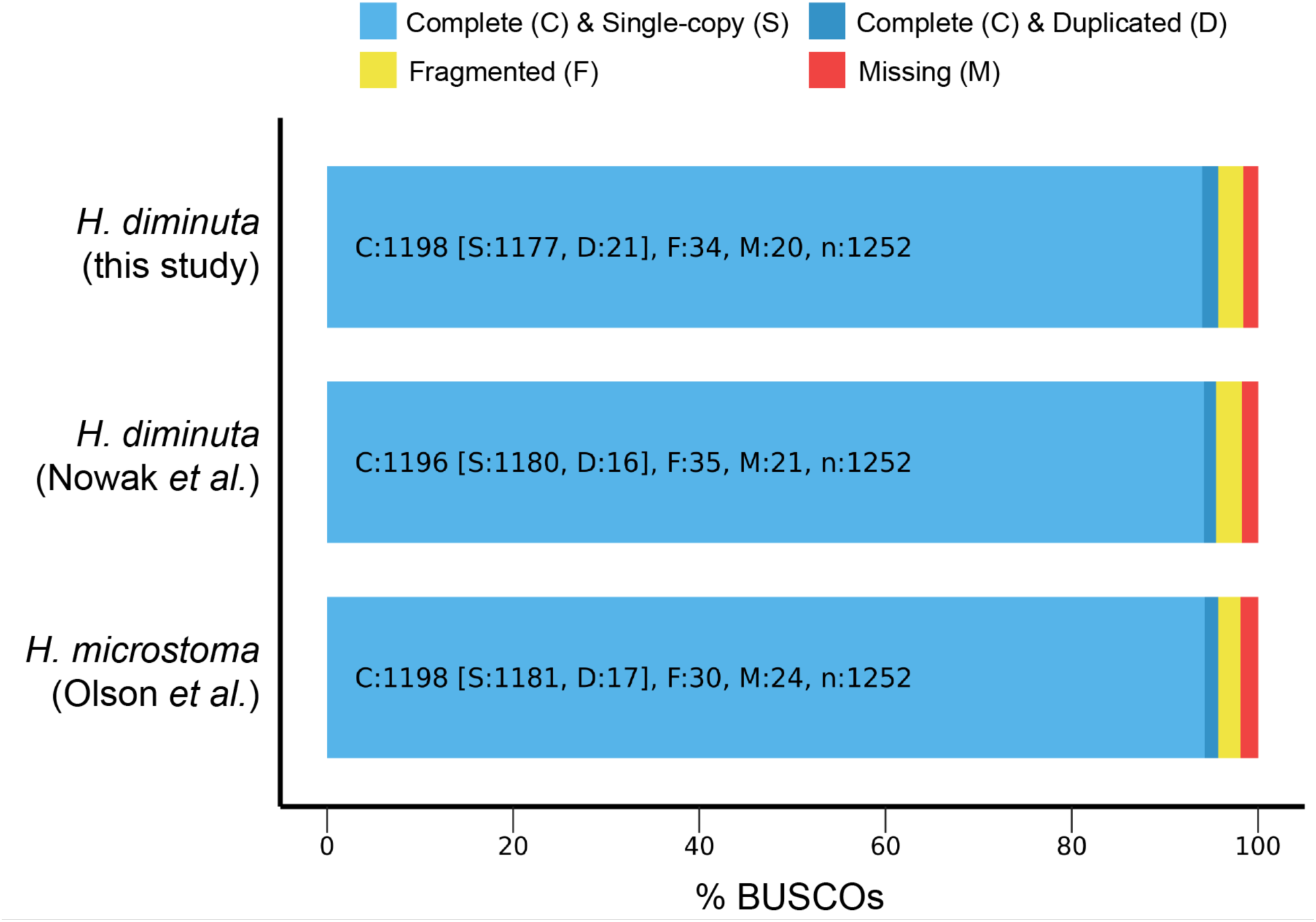
Estimated completeness of *Hymenolepis* genome assemblies. Bar charts show the estimated completeness of our chromosome-level assembly of *H. diminuta*, the *H. diminuta* assembly from Nowak et al. (2019), and the *H. microstoma* assembly from Olson et al. (2020) based on BUSCO analysis at the genome level using the Lophotrochozoan OrthoDB database.

### Hymenolepis genomes contain a high abundance of unclassified repeats

We inferred a library of interspersed repeats for *H. diminuta* using RepeatModeler applied to nuclear sequences in our chromosome-level assembly, then annotated and masked repeats using RepeatMasker. This analysis revealed that 28.1% of our *H. diminuta* chromosome-level assembly consists of interspersed repeats, with most of these sequences coming from currently unclassified repeat families (Figure 2, Supplementary Table 1). Of those repeats which can be classified, retroelement sequences (5.86%) occupy more than twice as much of the genome as DNA transposon sequences (2.16%). Repeat content estimated from annotation of the genome (28.1%) is higher, but on a similar order of magnitude, than that estimated from raw HiFi reads (21.8%) (Supplementary Figure 1). Both of our estimates of repeat content are much higher than what Nowak et al. (2019) previously reported for the *H. diminuta* genome (0.72%), who suggested that the low level of estimated repeat content they observed was caused by missing data in their assembly. To resolve this discrepancy, we annotated repeats in the *H. diminuta* assembly from Nowak et al. (2019) using the repeat library generated from our chromosome-level assembly. RepeatMasking the Nowak et al. (2019) assembly with our repeat library revealed a high overall abundance of interspersed repeats (22.9%) in their assembly with similar abundance trends for all categories of repeats as observed in our assembly (Figure 2). Hence, we conclude that the low repeat abundance for *H. diminuta* reported previously by Nowak et al. (2019) was not caused by missing data in their assembly, but rather by use of a non-hymenolepid repeat library.

**Figure 2.**
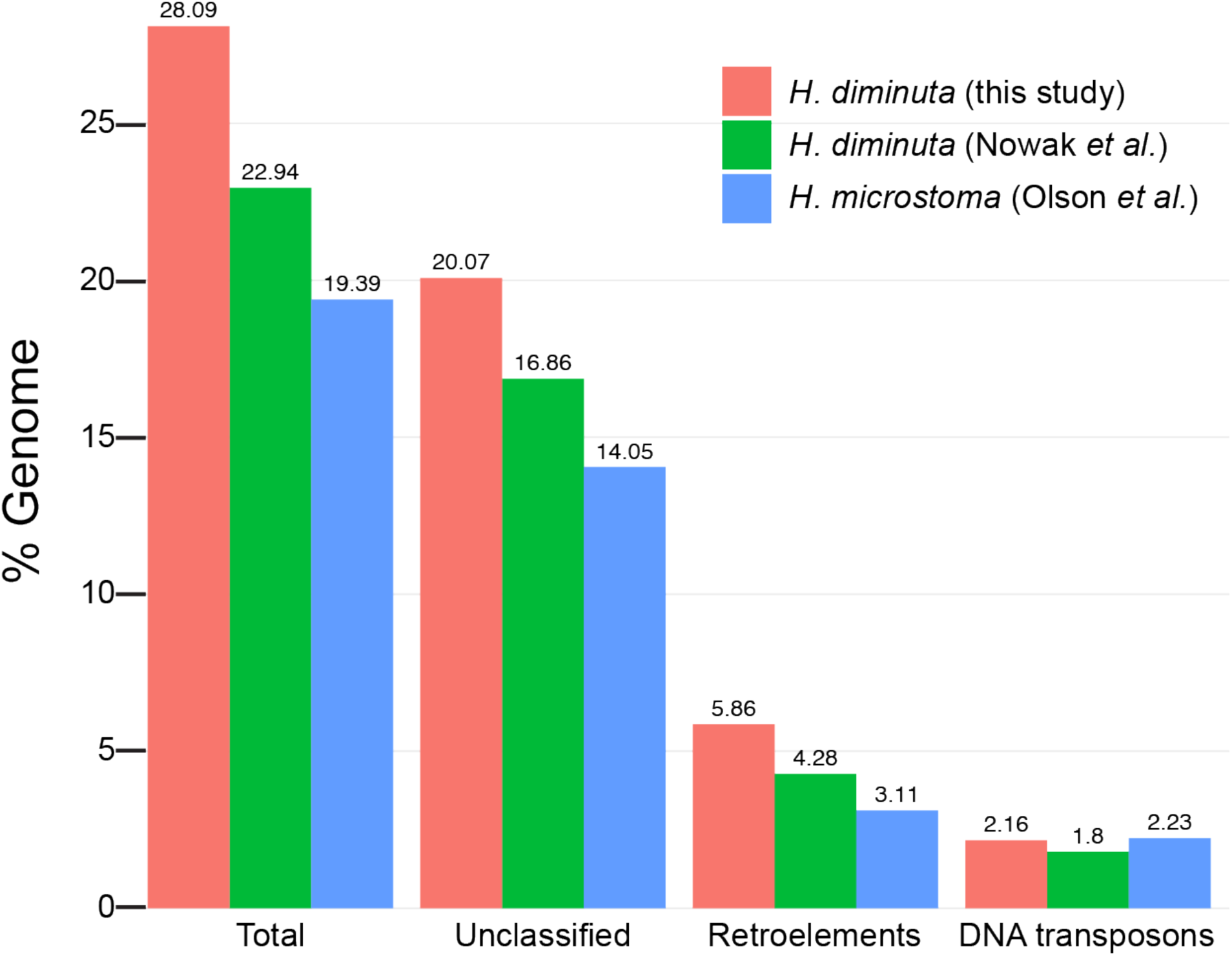
Abundance of interspersed repeats in *Hymenolepis* genomes. Bar charts show the proportions of selected major repeat types for our chromosome-level assembly of *H. diminuta*, the *H. diminuta* assembly from Nowak et al. (2019), and the *H. microstoma* assembly from Olson et al. (2020). Species-specific *de novo* repeat identification and annotation was performed using the *H. diminuta* assembly from this study and the *H. microstoma* assembly from Olson et al. (2020). To control for repeat library composition, the *de novo* repeat library from our *H. diminuta* genome was used to annotate repeats in the *H. diminuta* genome from Nowak et al. (2019).

To confirm whether the repeat landscape observed in *H. diminuta* is more broadly representative of *Hymenolepis* genomes in general, we applied the same repeat discovery and annotation procedure we used for our assembly to the *H. microstoma* assembly from Olson et al. (2020). We confirmed the previous findings by Olson et al. (2020) that a large proportion (19.4%) of the mouse bile-duct tapeworm genome consists of interspersed repeats, with qualitatively similar abundance trends for all categories of repeats as observed for our *H. diminuta* assembly (Figure 2). Relative to *H. diminuta*, a lower proportion of retroelement repeats but a similar proportion of DNA transposon repeats is observed in *H. microstoma*. This observation could be explained by inclusion of additional retroelement sequences in our *H. diminuta* assembly that are missing from the *H. microstoma* assembly or retroelement proliferation on the *H. diminuta* lineage since divergence from their common ancestor. Overall, we conclude that genomes for species in the genus *Hymenolepis* contain ∼20% interspersed repeats or more, with most repeat families and instances yet to be classified.

### Hymenolepis genomes contain approximately 10,000 protein coding genes

Previous studies on *Hymenolepis* genomes have reported the number of protein coding genes to range from ∼10,000 in *H. microstoma* (Olson et al. 2020) to ∼15,000 in *H. diminuta* (Nowak et al. 2019). This substantial difference in estimated gene number could be caused by variation in assembly quality, different gene prediction strategies, or extensive gene gain/loss among *Hymenolepis* species. To provide insight into this discrepancy and generate a genome annotation for our chromosome-level *H. diminuta* assembly, we predicted protein coding genes in our assembly, as well as in assemblies from Nowak et al. (2019) and Olson et al. (2020), using BRAKER3. Initial analysis of replicate BRAKER3 runs with different sources of evidence revealed that numbers of predicted protein coding genes and transcripts varied substantially for all assemblies as a function of the type of evidence used to train gene predictors (protein sequence homology, RNA-seq, or both), as well as more modest variation across replicate runs regardless of input evidence type (Figure 3). Regardless of input assembly, BRAKER3 runs that used both species-specific RNA-seq and protein sequence homology as evidence generally predicted the lowest number of genes (Figure 3A) but the highest number of transcripts (Figure 3B). Regardless of evidence type, BRAKER3 runs predicted the fewest gene models for the highly fragmented Nowak et al. (2019) assembly, and we never observed a BRAKER3 run for this assembly with more than 10,000 genes.

**Figure 3.**
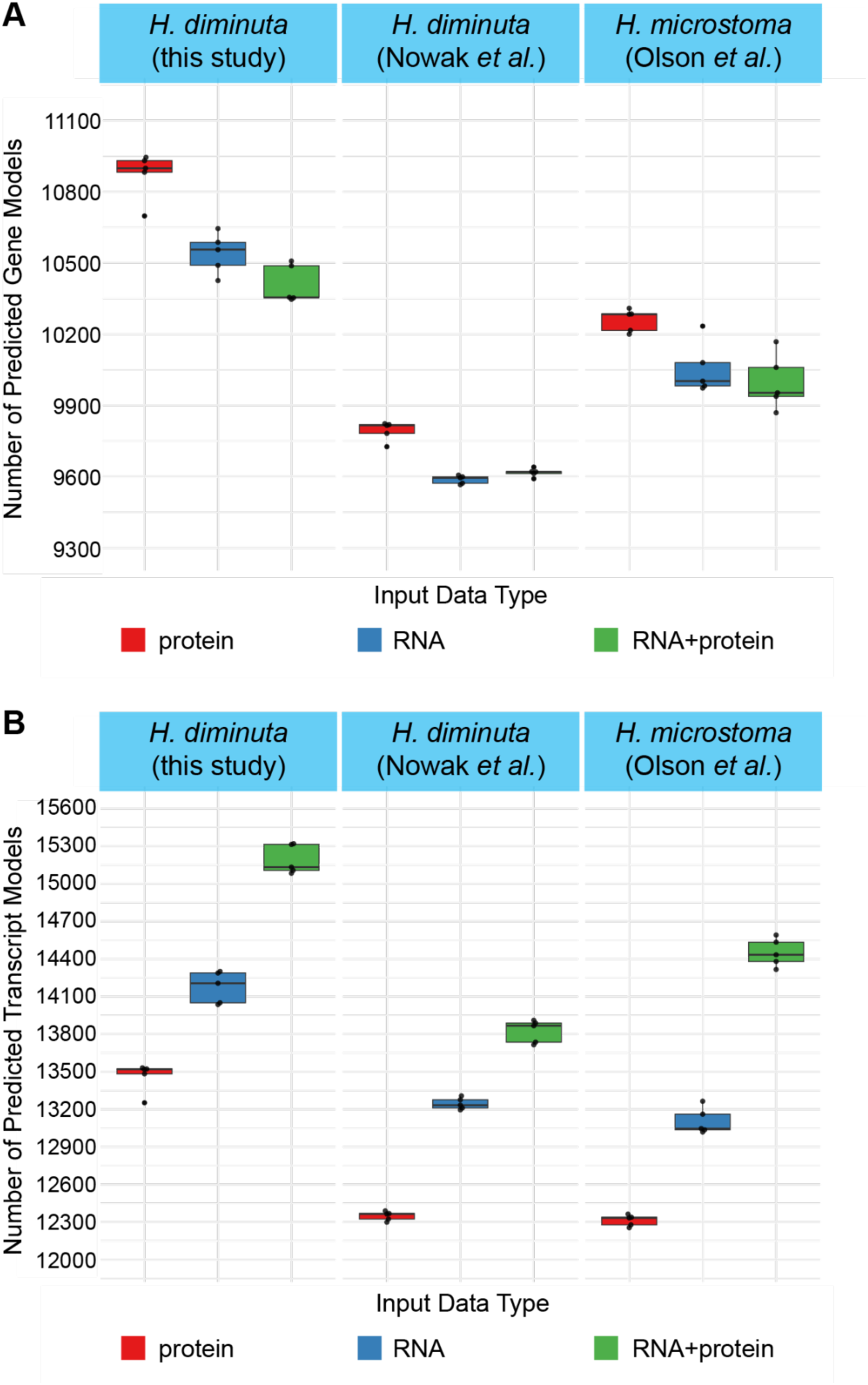
Variation in the number of protein coding genes and transcripts predicted by BRAKER3 across evidence type and replicates. Boxplots show the distribution of the number of (A) protein coding genes and (B) transcripts annotated by BRAKER3 across 5 replicate runs for our chromosome-level assembly of the *H. diminuta* genome, the *H. diminuta* assembly from Nowak et al. (2019), and the *H. microstoma* assembly from Olson et al. (2020), as a function of evidence type used to train gene predictors.

Based on this analysis, we selected a single BRAKER3 run that used both RNA-seq and protein evidence as the nuclear protein coding gene annotation for our *H. diminuta* chromosome-level assembly. To generate the final genome annotation, we merged this nuclear protein coding gene annotation with nuclear tRNA genes annotated using tRNAscan-SE, plus mtDNA protein coding and tRNA genes annotated using MitoFinder. Our final annotation contains 14,960 mRNA transcripts from 10,241 nuclear protein coding genes, 2,489 nuclear tRNAs, 36 mtDNA protein coding genes, and 22 mtDNA tRNAs (Table 2). 7,570 nuclear protein coding genes have at least one functional annotation predicted by EggNOG-mapper (Table 2). Using the longest isoform per gene to avoid artifactual inference of duplicated BUSCO genes, our annotation has a very high level of estimated completeness based on the high proportion of complete BUSCO genes (95.4%) present in the Lophotrochozoan OrthoDB database (Figure 4). BUSCO completeness estimated from our annotation (Figure 4) is only very slightly lower than BUSCO completeness estimated from our assembly (Figure 1), suggesting we have predicted at least one transcript from most protein coding genes present in our genome assembly.

**Figure 4.**
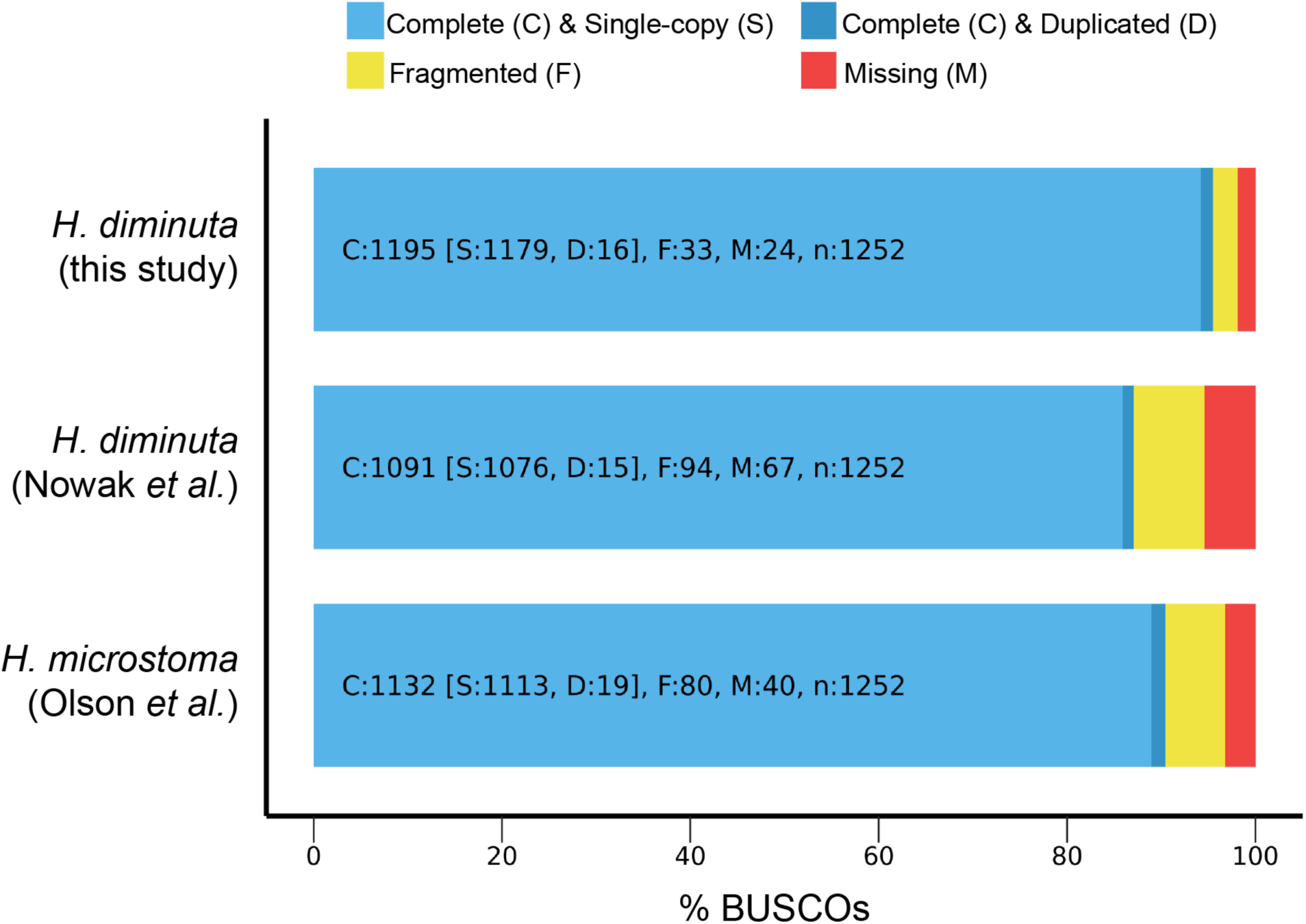
Estimated completeness of *Hymenolepis* genome annotations. Bar charts show the estimated completeness of the protein coding gene annotations for our chromosome-level assembly of *H. diminuta*, the *H. diminuta* assembly from Nowak et al. (2019), and the *H. microstoma* assembly from Olson et al. (2020) based on BUSCO analysis at the transcriptome level using the Lophotrochozoan OrthoDB database. In all cases, the longest isoform for each gene was selected for BUSCO analysis to prevent artifactual inference of duplicated genes caused by the presence of alternative isoforms.

**Table 2.** Annotation statistics for *Hymenolepis* genomes. Statistics shown are for BRAKER3 annotation of our chromosome-level assembly of *H. diminuta*, the *H. diminuta* annotation from Nowak et al. (2019) (Genbank: GCA_902177915.1) and the *H. microstoma* annotation from Olson et al. (2020) (Genbank: GCA_000469805.3).

|  | *H. diminuta*<br>(this study) | *H. diminuta*<br>(Nowak *et al.*) | *H. microstoma*<br>(Olson *et al.*) |
| --- | --- | --- | --- |
| Number of protein coding genes | 10,241 | 15,165 | 10,139 |
| Number of exons | 125,129 | 103,033 | 90,693 |
| Number of introns | 110,169 | 83,727 | 79,264 |
| Number of mRNAs | 14,960 | 19,306 | 11,429 |
| mean gene length (bp) | 7,566 | 4,260 | 6,883 |
| mean exon length (bp) | 216 | 230 | 218 |
| mean intron length (bp) | 1,092 | 850 | 868 |
| mean mRNA length (bp) | 1,809 | 1,049 | 1,729 |
| % of genome covered by genes | 41.5 | 36.5 | 41.3 |
| mean mRNAs per gene | 1.46 | 1.27 | 1.13 |
| mean exons per mRNA | 8.36 | 5.34 | 7.94 |
| mean introns per mRNA | 7.36 | 4.34 | 6.94 |
| Number of nuclear tRNA genes | 2,489 | N.A. | N.A. |
| Number of mtDNA protein coding genes | 36 | N.A. | N.A. |
| Number of mtDNA tRNA genes | 22 | N.A. | N.A. |

Relative to the published annotation from Nowak et al. (2019), our final genome annotation for *H. diminuta* predicts ∼5,000 fewer protein coding genes/mRNAs, predicts longer genes, covers a relatively larger percentage of the genome, has more exons and introns, more mRNAs per gene, and more exons/introns per mRNA (Table 2). Similar trends are observed when we filter annotation datasets to contain only the longest isoform per gene (Supplementary Table 2), with the exception that the Nowak et al. (2019) longest isoform dataset has more exons than our longest isoform dataset. Using longest isoform datasets to assess BUSCO completeness at the transcriptome level, we also observed many more fragmented and missing genes in the Nowak et al. (2019) annotation relative to our annotation (Figure 4). The Nowak et al. (2019) annotation also has a much higher rate of missing BUSCO genes (Figure 4) than estimated from their assembly (Figure 1), indicating that their annotation has many false negative gene predictions in addition to predicting ∼5,000 more genes than our annotation. To compare annotations on the same assembly coordinates, we lifted over the Nowak et al. (2019) longest isoform dataset to our chromosome-level assembly and compared annotation overlaps directly. This analysis revealed 4,602 genes from Nowak et al. (2019) with no evidence of a gene model in our annotation (putative false positive predictions in their annotation), and 1,315 genes in our annotation that had no corresponding gene model in the Nowak et al. (2019) annotation (putative false negative predictions in their annotation) (Supplementary Table 3). A large number of gene models in our annotation (n=813) were represented by more than one gene model in the Nowak et al. (2019) annotation (putative fragmented genes in their annotation), while very few Nowak et al. (2019) annotations (n=53) were represented by more than one gene model in our annotation (Supplementary Table 3).

Compared to the *H. microstoma* annotation reported by Olson et al. (2020), our annotation has a similar number of protein coding genes but >3,000 more mRNA transcripts, while the percentage of the genome assembly covered by genes is nearly the same in both species (Table 2). Additionally, our annotation has modestly longer gene and transcript lengths, more exons and introns, more mRNA isoforms per gene, and more exons/introns per mRNA. Longest isoform BUSCO analysis also reveals many fewer fragmented genes in our annotation relative to the *H. microstoma* annotation reported by Olson et al. (2020) (Figure 4). Finally, our annotation has fewer missing BUSCO genes than the *H. microstoma* gene annotation despite having an overall similar number of genes. These results suggest the *H. microstoma* annotation includes a proportion of false positive and false negative gene predictions on the same order as the false negative rate estimated by BUSCO (∼3%).

Overall, we conclude that *Hymenolepis* genomes have on the order of ∼10,000 protein coding genes and that the gene number of ∼15,000 for *H. diminuta* previously reported by Nowak et al. (2019) is an over-estimate that cannot be explained by run-to-run variation in BRAKER3, or the evidence type used to train BRAKER3 gene predictors. Our results indicate that the previous gene annotation for *H. diminuta* reported by Nowak et al. (2019) contains many potential false positive and false negative gene predictions, as well as many partial gene models that have been merged into fewer, more complete gene models in our annotation of the *H. diminuta* genome. Our annotation also shows improved protein coding transcript structure and diversity relative to the *H. microstoma* annotation reported by Olson et al. (2020). Finally, we note that our annotation of the *H. diminuta* genome also includes nuclear tRNAs, mtDNA protein coding genes, and mtDNA tRNAs, none of which are represented in the previous genome annotations for *H. diminuta* or *H. microstoma* (Table 2), making it the most comprehensive *Hymenolepis* genome annotation to date.

### Major features of genome architecture are conserved among Hymenolepis genomes

Previous work has shown that *H. microstoma* genes are found in conserved syntenic regions that are broadly conserved with distantly related helminths, with extensive shuffling of gene order within these syntenic regions (Olson et al. 2020). To understand the pattern of gene order evolution at a shorter timescale within the *Hymenolepis* genus, we identified orthogroups using our *H. diminuta* annotation and the *H. microstoma* annotation from Olson et al. (2020) and visualized patterns of gene order evolution in GENESPACE. As shown in Figure 5, gene order in the *Hymenolepis* genus is broadly conserved within all nuclear chromosomes. However, we detect 65 disruptions in gene order due to putative inverted genome segments of chromosomes in the two species. Pairwise alignment of our *H. diminuta* assembly to the *H. microstoma* assembly from Olson et al. (2020) confirms that all large-scale disruptions in gene order observed are caused by inversions present on all six chromosomes (Supplementary Figure 2).

**Figure 5.**
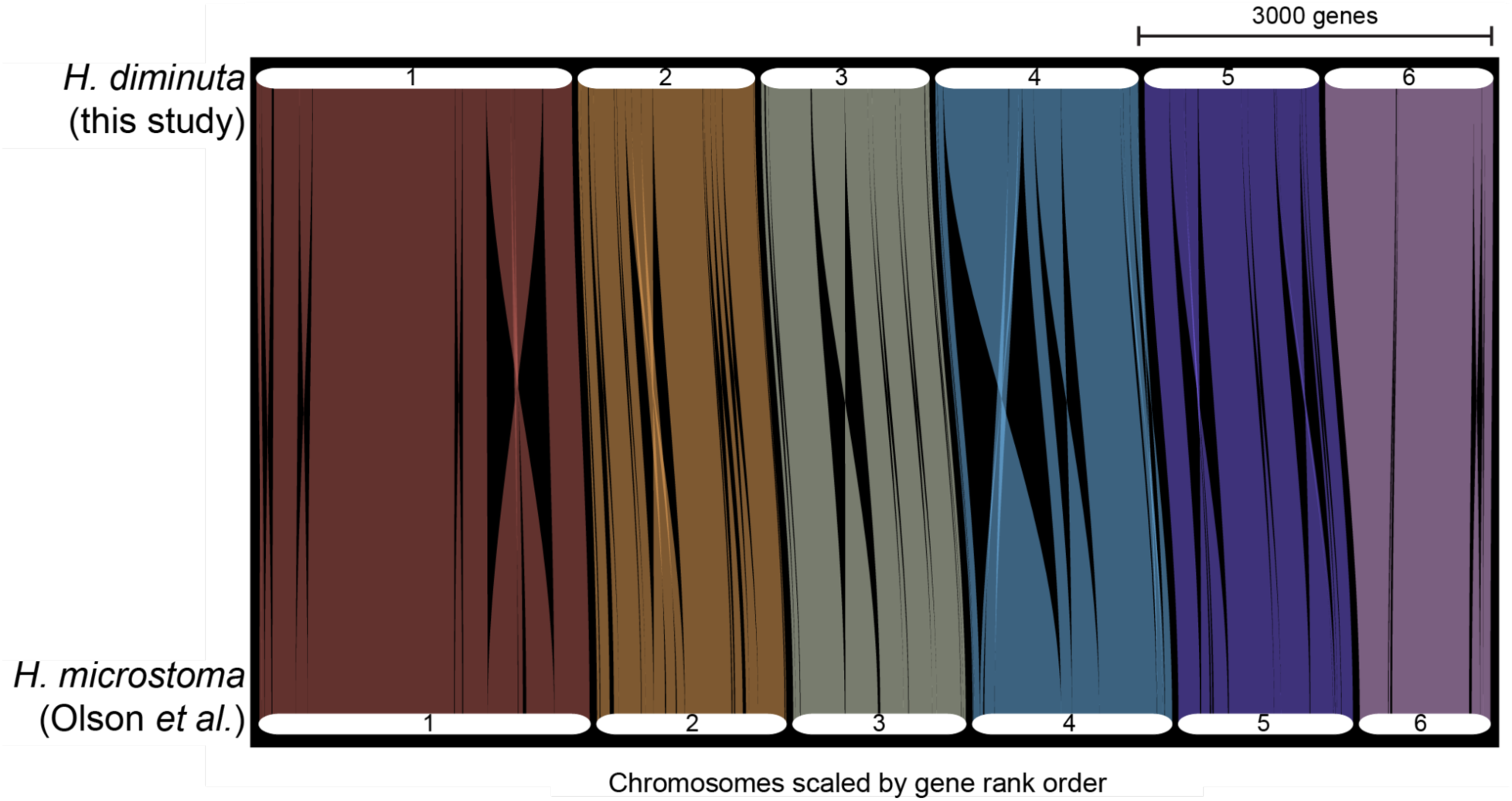
Evolution of gene order between *H. diminuta* and *H. microstoma.* GENESPACE riparian plot comparing the order of conserved genes between *H. diminuta* (top) and *H. microstoma* (bottom). Each ribbon represents a syntenic block of orthogroups, inferred by OrthoFinder using the longest isoform per gene, with chromosome lengths scaled by the number of genes. Crossing ribbons within chromosomes indicate putative intrachromosomal inversions.

To verify that putative inversion breakpoints were not caused by assembly artifacts, we mapped PacBio raw reads from *H. diminuta* and *H. microstoma* back to their respective genomes and visualized patterns of high-quality, unique read mappings in putative breakpoint regions. We observed direct evidence for contiguous overlapping mapped reads supporting proper assembly of 58/65 putative inversion breakpoint regions in *H. diminuta* (e.g., Supplemental Figure 3A). The remaining 7 putative inversion breakpoint regions contain highly repetitive sequences that prevent high-quality, unique read mapping support (e.g., Supplemental Figure 3B), assembly gaps that disrupt read mapping contiguity (e.g., Supplemental Figure 3C), or both. We emphasize that these latter two categories provide evidence neither for nor against misassembly in putative breakpoint regions. Similarly, we observed direct evidence for contiguous overlapping read support for 57/64 putative inversion breakpoint regions in *H. microstoma*, with the remaining 7 putative inversion breakpoint regions containing highly repetitive sequences, assembly gaps, or both. We note that one fewer putative inversion breakpoint region was observed in *H. microstoma* than *H. diminuta* because of a complex rearrangement that reuses the first breakpoint region on chromosome 1. 49 inversion breakpoint regions are well supported in both species, providing direct evidence that most disruptions in gene order observed between *H. diminuta* and *H. microstoma* are caused by *bona fide* genome rearrangements rather than misassembly in one or both genomes. Overall, we conclude that the gene content of chromosomes in the genus *Hymenolepis* is broadly conserved and that the vast majority of disruptions in gene order between the *H. diminuta* and *H. microstoma* genomes is due to at least 49 intrachromosomal inversions spread across all chromosomes.

Lastly, we sought to determine whether *H. diminuta* chromosome ends have the same unusual organization as observed in *H. microstoma*, with one chromosomal end terminating in telomeric repeats and the other end terminating in centromeric repeats (Olson et al. 2020). To do this, we predicted telomeric and centromeric repeats in our chromosome-level assembly using tidk and centroAnno, respectively. We identified canonical telomeric repeats at one end of all 6 chromosomes, and centromeric repeats at the other end of all chromosomes in our assembly except for chromosome 5 (Figure 6). We find no evidence that the largest *H. diminuta* chromosome has a centromere repeat array in a sub-metacentric location as suggested by cytogenic data in one (Mutafova and Gergova 1994) but not another (Kisner 1957) previous study. Telomeric and centromeric ends in *H. diminuta* are in the same relative orientation as observed in *H. microstoma* (Olson et al. 2020). These results indicate that our chromosome assembly is “telomere-to-centromere” complete for 5 chromosomes and that the unusual organization of chromosomal termini first identified in *H. microstoma* (Olson et al. 2020) is likely to be conserved in the genus *Hymenolepis*. To investigate the alternative hypothesis that the unusual pattern of chromosome ends terminating in centromere repeats at one end might be due to incomplete assembly at centromeric ends, we also predicted telomeric and centromeric repeats in unplaced contigs (Supplemental Figure 4). We identified two unplaced contigs over 1 Mb that contained both telomeric and centromeric repeats (ptig000012l and ptig000019l) that could potentially represent extensions to centromeric ends with conventional telomeres. However, in both cases the telomeric repeats are not arranged in tandem arrays as expected for *bona fide* telomeres. Thus, we find no evidence to support the hypothesis that *Hymenolepis* genomes have conventional chromosomes capped by telomeres on both ends, as found in other non-hymenolepid tapeworms (Tsai et al. 2013; Li et al. 2018).

**Figure 6.**
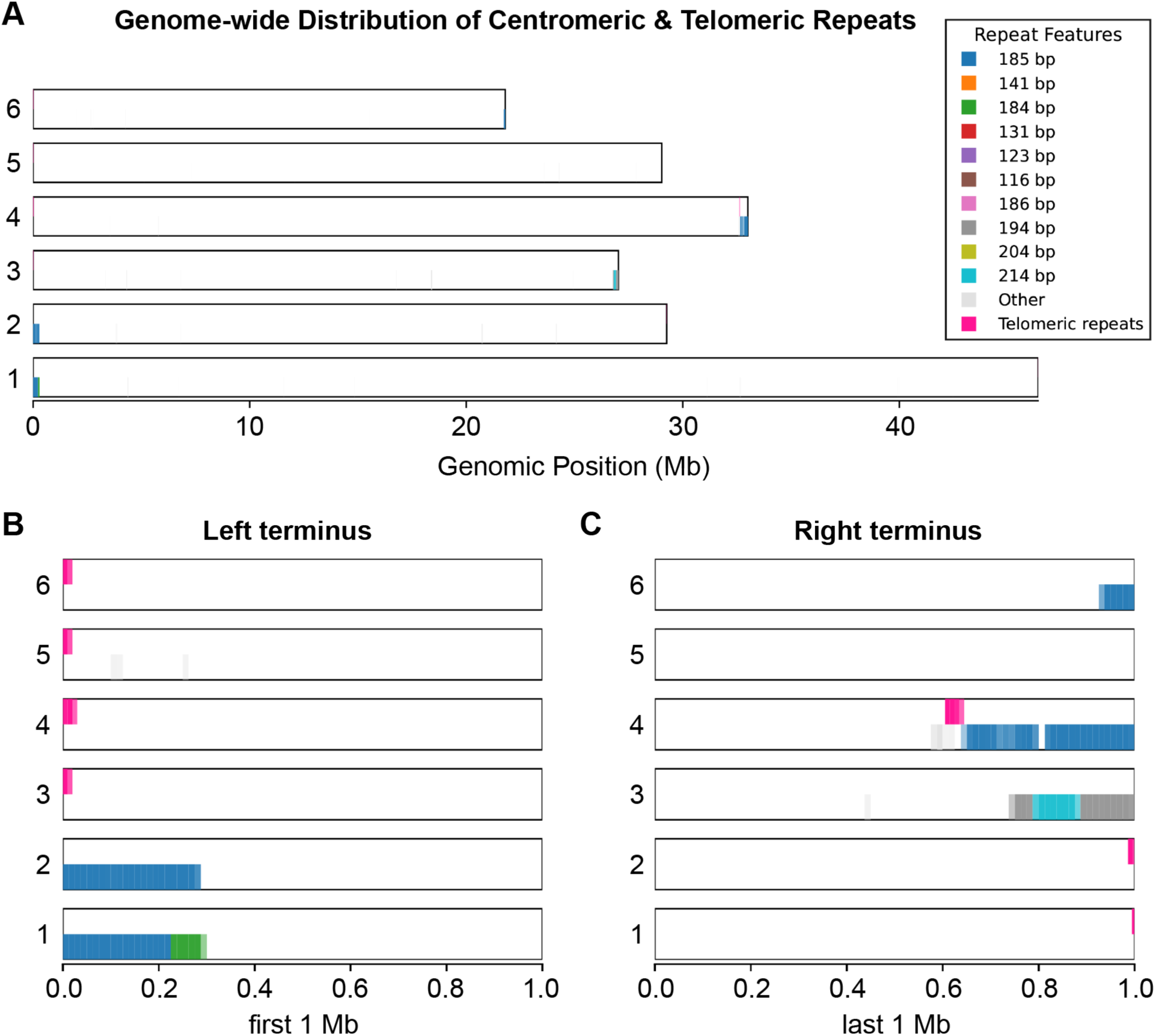
Distribution of centromeric and telomeric repeats across the *H. diminuta* genome. (A) Genomic distribution of potential centromeric repeats (identified with centroAnno) and canonical telomeric repeats (identified with Tidk) across each of the six *H. diminuta* chromosomes, plotted by genomic position (Mb). Centromeric repeats are colored by the most frequent monomer length classes (with remaining monomers grouped as “Other”) and telomeric repeats are shown in magenta. (B,C) Zoomed views of the first 1 Mb (left terminus, B) and last 1 Mb (right terminus, C) of each chromosome, showing the localization of predicted telomeric and centromeric repeats near the termini of each chromosome.

## Conclusions

Here we used PacBio HiFi sequencing and assembly to generate a high-quality genome assembly and annotation for *H. diminuta*. This resource will accelerate molecular studies and open new possibilities for genome-level analyses that will greatly improve the utility of this important model organism. This resource will enhance the utility of *H. diminuta* as a model for studying tapeworm growth, regeneration, and reproduction. Our assembly also supports the finding that *Hymenolepis* chromosomes have one telomeric and one centromeric end inviting further study of this unusual phenomenon.

## Data Availability Statement

Raw HiFi sequencing reads are available in NCBI Sequence Read Archive (SRR33683259) under NCBI BioProject PRJNA1257364. The genome assembly and associated annotations are available under NCBI Genome accession GCA_056153285.1. All custom code and scripts used for the data analyses in this paper are available on Zenodo (https://doi.org/10.5281/zenodo.20737624).

## Acknowledgements

We thank the Arizona Genomics Institute, Tucson, AZ for services including DNA extraction, PacBio library preparation, and HiFi sequencing; the Georgia Advanced Computing Resource Center at the University of Georgia for technical support and computational resources. We used Claude for drafting and debugging selected scripts used for analysis. All AI-assisted code was reviewed, tested, modified as needed, and validated by the authors.

## Supplementary Figures

**Supplementary Figure 1.**
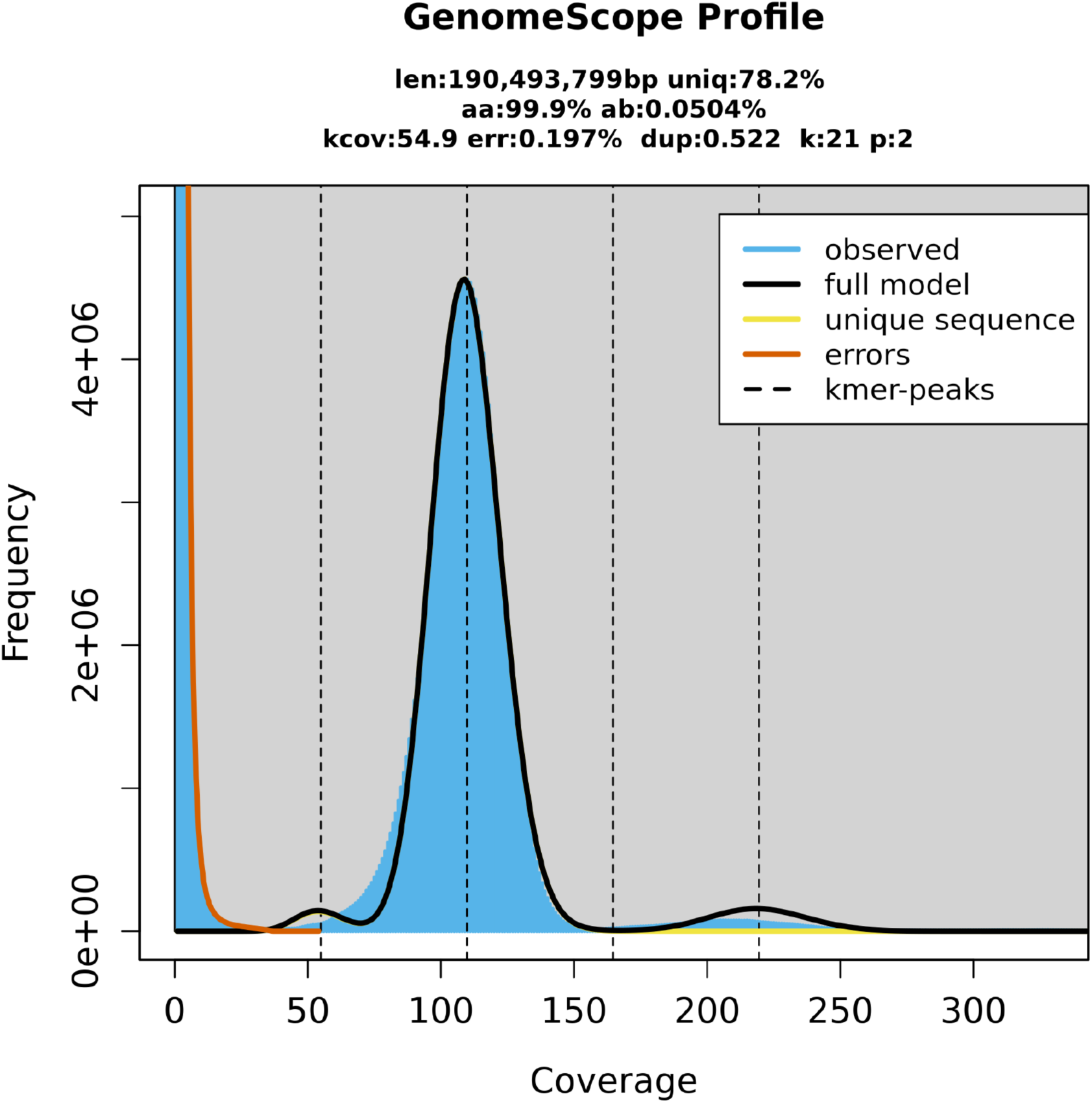
Estimates of genome size, fold-coverage, repetitiveness, and heterozygosity based on unassembled PacBio HiFi reads for *H. diminuta*. “ab” represents the level of heterozygosity, and “kcov” represents the coverage of heterozygous kmers. 2*kcov provides an estimate of the fold coverage for homozygous regions of the genome.

**Supplementary Figure 2.**
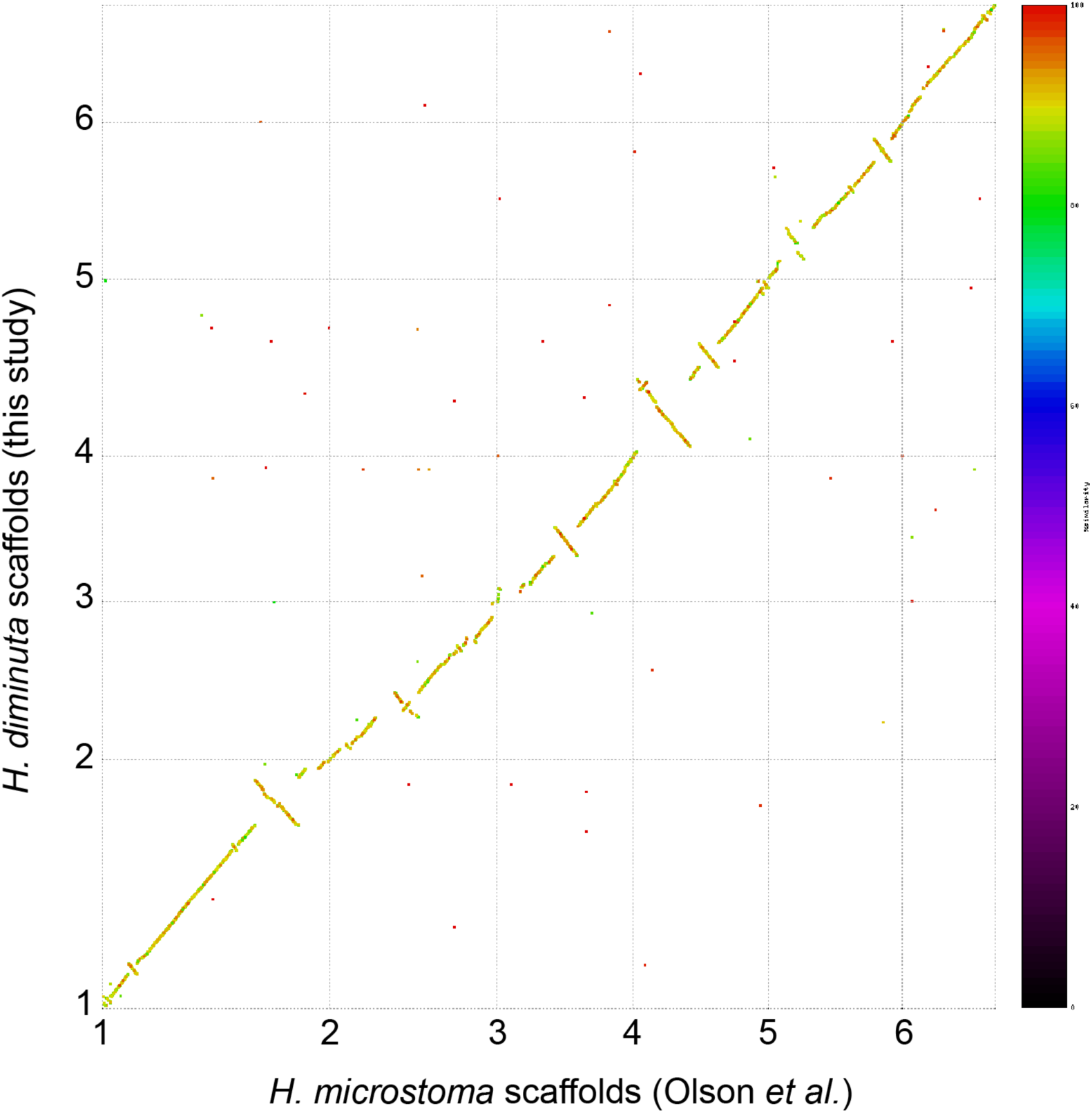
Pairwise alignment of *H. diminuta* and *H. microstoma* genomes. Shown is a Mummerplot of nuclear chromosomal scaffolds for our chromosome-level assembly of *H. diminuta* on the Y-axis and the *H. microstoma* assembly from Olson et al. (2020) on the X-axis. Thin dashed lines represent the boundaries of chromosomal assemblies. Dots represent homologous segments colored by the level of sequence similarity shown in the scale bar.

**Supplementary Figure 3.**
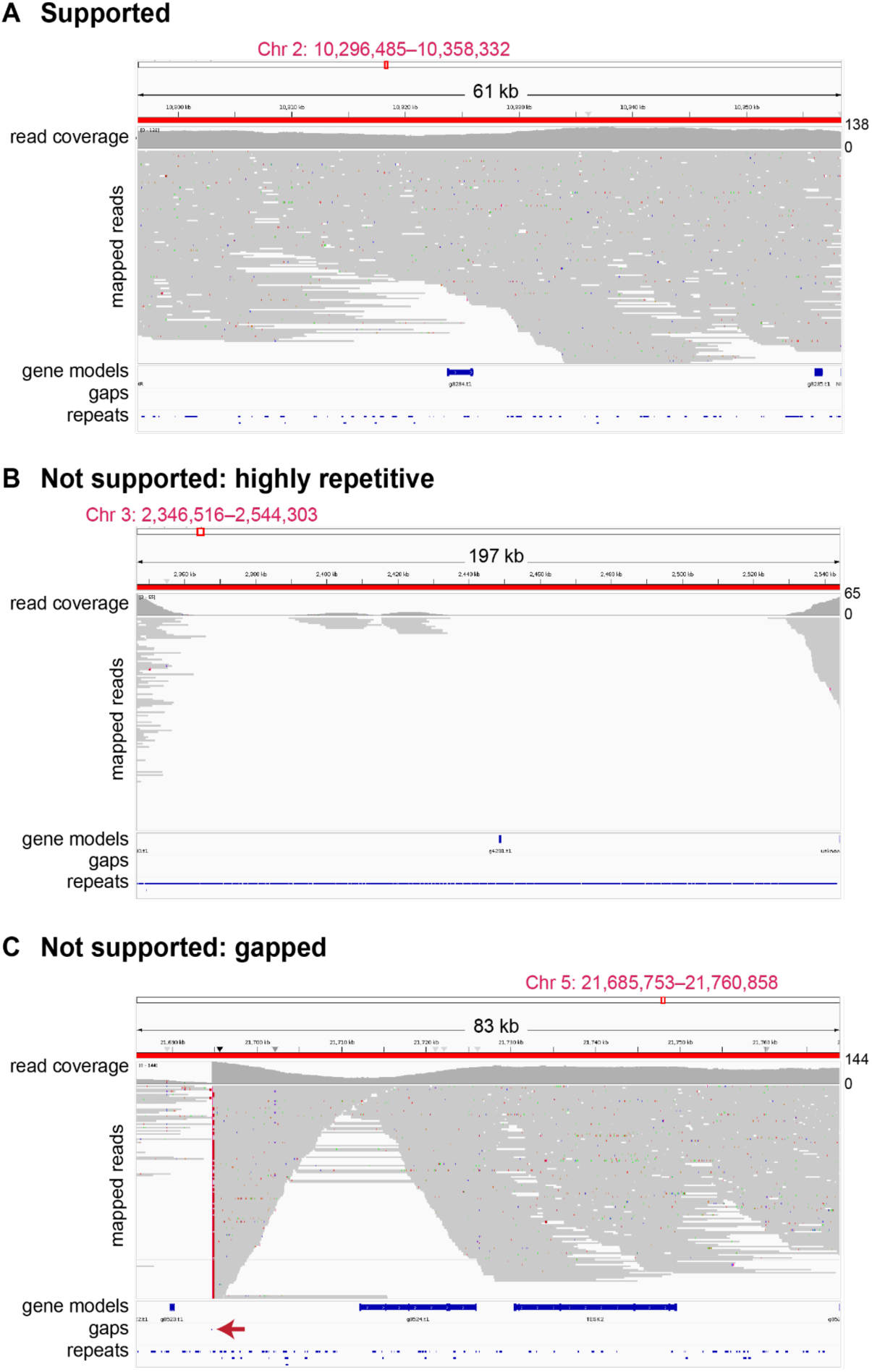
Examples of patterns of read support in putative inversion breakpoint regions identified by GENESPACE. Putative inversion breakpoint regions in our assembly of *H. diminuta* and the *H. microstoma* from Olson et al. (2020) were visualized in IGV in the context of high-quality (MAP ≥ 30), uniquely mapped reads together with assembly gene, repeat, and gap annotation tracks. Putative breakpoint regions were classified into three categories as show in the following examples from *H. diminuta*: A) Putative breakpoint region supported by contiguous overlapping high-quality uniquely mapped reads; B) Putative breakpoint region not supported by contiguous overlapping high-quality mapped reads due to high repeat content; C) Putative breakpoint region not supported by contiguous overlapping high-quality mapped reads due to assembly gap. We note that categories B and C are not mutually exclusive.

**Supplementary Figure 4.**
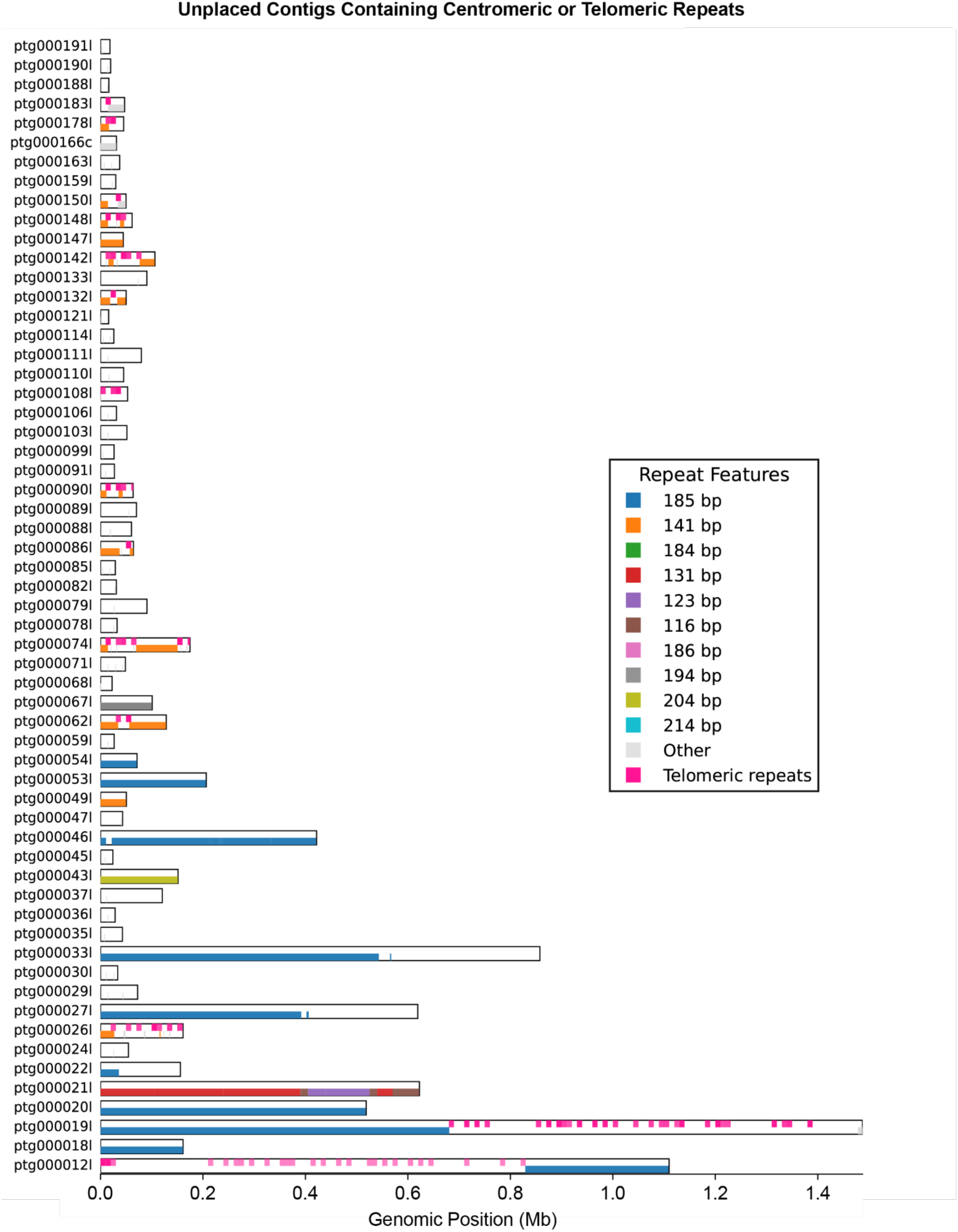
Distribution of centromeric and telomeric repeats across unplaced contigs of *H. diminuta* assembly. Genomic distribution of potential centromeric repeats (identified with centroAnno) and canonical telomeric repeats (identified with Tidk) along unplaced contigs, plotted against genomic position (Mb). Centromeric repeats are colored by monomer length, and telomeric repeats are shown in magenta.

## Supplementary Tables

**Supplementary Table 1.** Abundance of interspersed repeats in *Hymenolepis* genomes. Numbers of elements, total number of base pairs covered, and percent genome annotated by major repeat types for our chromosome-level assembly of *H. diminuta* (this study), the *H. diminuta* genome from Nowak et al. (2019), and the *H. microstoma* genome from Olson et al. (2020). Species-specific *de novo* repeat identification and annotation was performed using the *H. diminuta* assembly from this study and the *H. microstoma* assembly from Olson et al. (2020). To control for repeat library composition, the *de novo* repeat library from our *H. diminuta* genome was used to annotate repeats in the *H. diminuta* genome from Nowak et al. (2019).

| Repeat Type | Metric | *H. diminuta*<br>(this study) | *H. diminuta*<br>(Nowak *et al.*) | *H. microstoma*<br>(Olson *et al.*) |
| --- | --- | --- | --- | --- |
| Total interspersed repeat | Number of elements | 200,626 | 184,275 | 108,505 |
|  | Coverage (bp) | 52,400,259 | 40,617,056 | 32,756,615 |
|  | % genome | 28.09 | 22.49 | 19.39 |
| Unclassified | Number of elements | 163,528 | 151,827 | 84,362 |
|  | Coverage (bp) | 37,437,857 | 29,849,755 | 23,732,282 |
|  | % genome | 20.07 | 16.86 | 14.05 |
| Retroelements | Number of elements | 29,981 | 25,697 | 14,374 |
|  | Coverage (bp) | 10,929,791 | 7,573,368 | 5,254,473 |
|  | % genome | 5.86 | 4.28 | 3.11 |
| DNA transposons | Number of elements | 7,117 | 6,751 | 9,769 |
|  | Coverage (bp) | 4,032,611 | 3,193,933 | 3,769,860 |
|  | % genome | 2.16 | 1.80 | 2.23 |

**Supplementary Table 2.** Annotation statistics for the longest isoform annotated in *Hymenolepis* genomes. Statistics shown are for the longest isoform per gene from the BRAKER3 annotation of our chromosome-level assembly of *H. diminuta*, the *H. diminuta* assembly from Nowak et al. (2019) (Genbank: GCA_902177915.1) and the *H. microstoma* assembly from Olson et al. (2020) (Genbank: GCA_000469805.3).

|  | *H. diminuta*<br>(this study) | *H. diminuta*<br>(Nowak *et al.*) | *H. microstoma*<br>(Olson *et al.*) |
| --- | --- | --- | --- |
| Number of protein coding genes | 10,241 | 15,165 | 10,139 |
| Number of exons | 69,291 | 70,453 | 73,590 |
| Number of introns | 59,050 | 55,288 | 63,451 |
| Number of mRNAs | 10,241 | 15,165 | 10,139 |
| mean gene length (bp) | 7,566 | 4,259 | 6,883 |
| mean exon length (bp) | 232 | 236 | 225 |
| mean intron length (bp) | 1,019 | 835 | 838 |
| mean mRNA length (bp) | 7,432 | 4,132 | 6,871 |
| % of genome covered by genes | 41.5 | 36.5 | 41.3 |
| mean mRNAs per gene | 1.00 | 1.00 | 1.00 |
| mean exons per mRNA | 6.77 | 4.65 | 7.26 |
| mean introns per mRNA | 5.77 | 3.65 | 6.26 |
| Number of nuclear tRNA genes | 2489 | N.A. | N.A. |
| Number of mtDNA protein coding genes | 36 | N.A. | N.A. |
| Number of mtDNA tRNA genes | 22 | N.A. | N.A. |

**Supplementary Table 3.** Overlap of gene models from this study relative to gene models from Nowak et al. (2019). The longest isoform annotation dataset from the *H. diminuta* assembly of Nowak et al. (2019) was lifted over to our *H. diminuta* assembly and overlapped with the longest isoform annotation dataset from the current study to assess gene annotation correspondence in the two studies.

| Number of genes<br>from this study | Number of lifted-over<br>genes from Nowak *et al.* | Number of<br>cases |
| --- | --- | --- |
| 0 | 1 | 4602 |
| 1 | 0 | 1315 |
| 1 | 1 | 8005 |
| 1 | 2 | 541 |
| 1 | 3 | 148 |
| 1 | 4 | 64 |
| 1 | 5 | 22 |
| 1 | 6 | 11 |
| 1 | 7 | 12 |
| 1 | 8 | 9 |
| 1 | 9 | 3 |
| 1 | 11 | 1 |
| 1 | 12 | 2 |
| 2 | 1 | 51 |
| 3 | 1 | 2 |

## Additional Data Files on Zenodo

**Additional Data File 1.** RepeatModeler fasta file

**Additional Data File 2.** Genespace synteny file

**Additional Data File 3.** Breakpoint file

**Additional Data File 4.** Centromere repeat fasta file

**Additional Data File 5.** Centromere repeat annotation file

**Additional Data File 6.** Telomere repeat annotation file

**Additional Data File 7.** Zip file of code used for this study.

